# Lupin (*Lupinus sp.*) seeds exert anthelmintic activity associated with their alkaloid content

**DOI:** 10.1101/530956

**Authors:** O. Dubois, C. Allanic, C.L. Charvet, F. Guégnard, H. Février, I. Théry-Koné, J. Cortet, C. Koch, F. Bouvier, T. Fassier, D. Marcon, J.B. Magnin-Robert, N. Peineau, E. Courtot, C. Huau, A. Meynadier, C. Enguehard-Gueiffier, C. Neveu, L. Boudesocque-Delaye, G. Sallé

## Abstract

The growing expansion range of drug resistant parasitic nematode populations threatens the sustainability of ruminant farming worldwide. In this context, nutraceuticals, *i.e.* feed that would both fulfil dietary requirements while ensuring parasite control, would contribute to increase farming sustainability in developed and low resource settings. In this study, we characterized the anthelmintic potential of lupin seed extracts against major ruminant trichostrongylids, *i.e. H. contortus* and *T. circumcincta*. Our observations showed that total seed extracts from commercially available lupin varieties significantly inhibited larval migration. This anthelmintic effect was sustained on multidrug resistant field isolate and across parasite species and was mediated by the seed alkaloid content. Analytical chemistry revealed a set of four lupanine and sparteine-derivatives with anthelmintic activity and electrophysiology assays on recombinant nematode acetylcholine receptors suggested an antagonistic mode of action for lupin alkaloids. While commercial lupin seeds did not exert direct anthelmintic effect in *H. contortus* infected lupin-fed ewes and goats, it significantly dampened blood production losses suffered by goats. Lupin seed extracts hence provide a working basis for the development of novel anthelmintic compounds able to break drug resistance in the field, while they could be used to increase the resilience of infected dairy ruminants.

## Introduction

Anthelmintic resistance is a major issue for the sustainable management of both human-^1^ and livestock-infective helminth species^2-4^. Gastro-intestinal parasitic nematodes (GIN) significantly impact human development, amounting to a loss of 10 million disability-adjusted life years^5^ and impeding the livestock industry with production losses^6,7^. While novel control solutions are urgently needed, only few anthelmintic compounds have been released in the recent years, like monepantel^8^, for which field drug resistance has been described only a couple of years after its release^9^. Vaccine remain difficult to design^10^ and GIN show enough transcriptomic plasticity to circumvent the vaccinal response of their host^11^. In the veterinary setting, the breeding of more resistant individuals in less need for treatment^12,13^, the implementation of targeted-selected treatment approaches^14^, or the use of tannin-rich plant extracts^15,16^ have been studied. This latter strategy relies on the combined properties of forages that show both good nutritional properties and bioactive compounds, known as “neutraceuticals”^15^. This approach should limit drug residues, lower drug selection pressure and can be deployed under low resource settings^16^.

Lupin (*Lupinus sp.)* is a grain legume that belongs to the genistoid clade of the Fabaceae family^17^. It produces protein- and energy-rich seeds used for ruminants^18^ or laying hens^19^ feeding, and contributes to reduce risk of obesity, diabetes and cardiovascular diseases in humans^20^.

Lupin also exerts antifungal properties^21^ and was shown to lower soil burden of the vine parasitic nematode *Xiphinema index*^22^. These antiparasitic properties are thought to be mediated by lupin quinolizidine and piperidine alkaloid compounds, that confer both bitterness and toxicity to the alkaloid-rich lupin varieties^23^. Indeed, alkaloidic compounds, like lupanine and spartein, block the excitatory neuro-transmission through the binding of nicotinic acetylcholine receptors, nAChRs^23,24^. Interestingly, nAChRs are well characterized pharmacological targets for the control of parasitic nematodes^25^. These transmembrane ligand-gated ion channels are made of five subunits that associate together to form homo- or heteropentameric receptors^25^. Widely used anthelmintics *i.e.* levamisole^26^, pyrantel^27^, and monepantel^28^, are agonists of these receptors, whereas derquantel, a derivative from the oxindole alkaloid paraherquamide^29^, acts as a nAChR antagonist^30^.

Therefore, lupin seed could harbour novel anthelmintic agents of great interest for GIN control and may be used as a nutraceutical for grazing ruminants. In this study, we investigated the anthelmintic potential of lupin seed by exposing major parasitic trichostrongylids, *i.e. Haemonchus contortus* and *Teladorsagia circumcincta*, to lupin seed extracts revealing its anthelmintic potential. We performed analytical chemistry to unravel four alkaloid compounds underpinning this effect and electrophysiology assay demonstrated their antagonist mode of action against nematode acetylcholine receptors. *In vivo* trial with commercial lupin seed in growing ewe and goats was also implemented.

## Results

### Lupin seed extracts show anthelmintic effect on *H. contortus* infective larvae

As a first step, aqueous extracts from 11 alkaloid-rich and -poor lupin seeds (Supplementary Table 1) were tested against drug-susceptible and multidrug-resistant *H. contortus* infective larvae using a larval migration assay (Fig. 1, Supplementary Table 2). Every aqueous extract considered (but LANG172) exerted a significant reduction of larval migration across parasite isolates whatever its anthelmintic resistance status (Fig. 1, Supplementary Table 2). Extracts from alkaloid-rich seeds demonstrated a 77.7% inhibition of larval migration and were thus generally more potent than the alkaloid-poor varieties (27.1% ± 0.04% difference in average inhibition, *F*_*1,63*_ = 28.68, *P*<10^−4^; Fig. 1, Supplementary Table 2). Among alkaloid-poor varieties, extracts from ENERGY and LL049 exerted similar potencies across parasite isolates (57.8% ± 0.2% and 59.3% ± 0.16% respectively) that significantly out-performed other alkaloid-poor varieties inhibitory effects (16.8% ± 0.03% difference, *F*_*2,27*_ = 62.8, *P* <10^−4^, Fig. 1).

**Figure 1.**
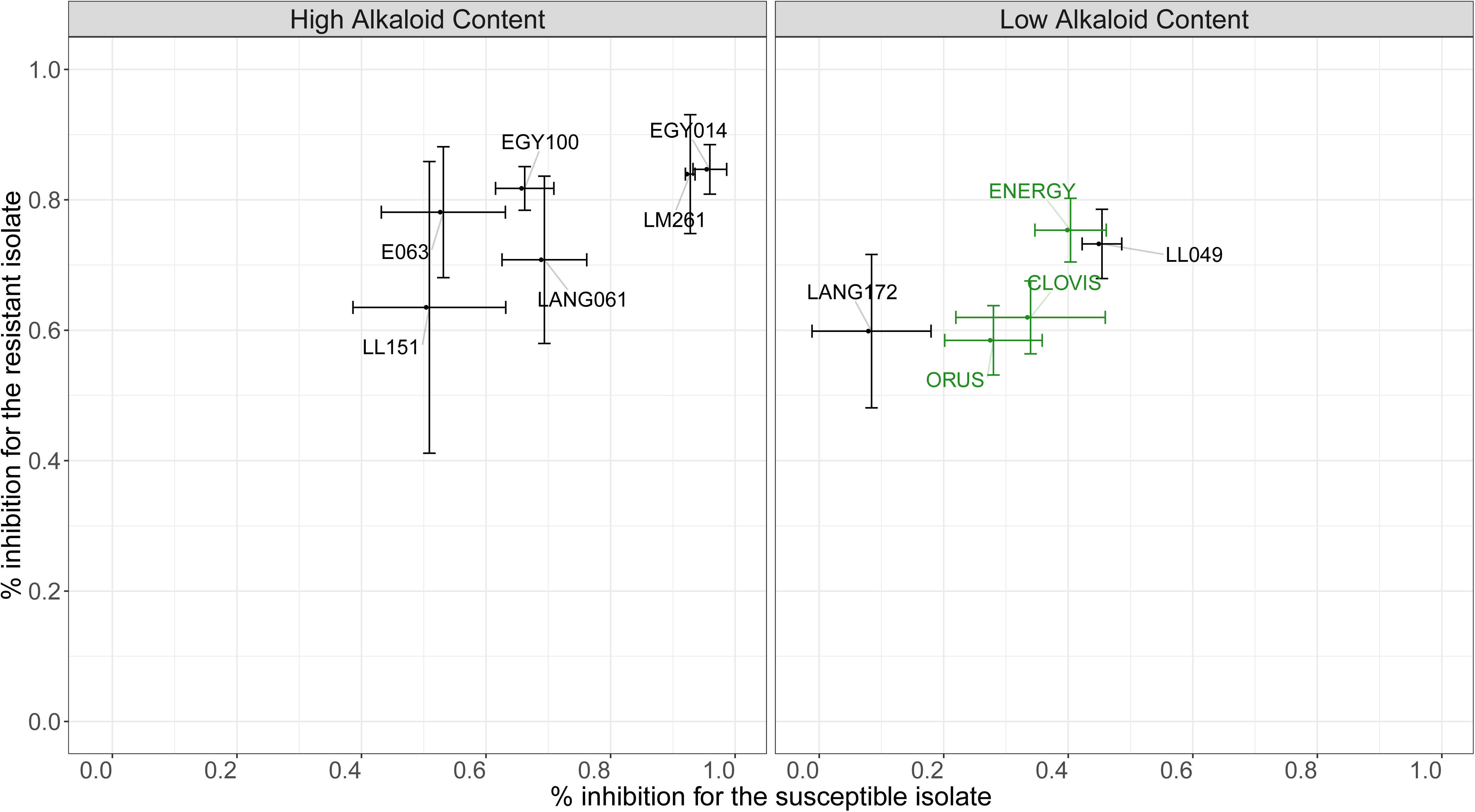
Initial screening highlights ENERGY and E063 as the most potent varieties. Figure depicts the average percentage of larval migration inhibition (relative to negative control) measured on drug-susceptible (x-axis) and drug-resistant (benzimidazole, levamisole, ivermectin; y-axis) *Haemonchus contortus* isolates exposed to 11 lupin varieties seed total extracts. Lines show standard deviation measured from three replicates. Green dots stand suitable for commercially available varieties.

In an initial attempt to establish the contribution of alkaloids to the inhibitory effect, alkaloids were extracted from alkaloid-rich varieties to evaluate their inhibitory effect. Alkaloid fractions inhibitory effects (Supplementary Table 3) were minimal for LL151 on the fully susceptible *H. contortus* isolate (52% ± 2.77% s.d.) and the most potent effect was observed for E063 on the multidrug resistant *H. contortus* isolate (82% ± 3.58% s.d.).

After this initial screening, the anthelmintic potential of lupin seed extracts was demonstrated against both drug-susceptible and multidrug-resistant isolate. One alkaloid-rich and alkaloid-poor were selected for further study. The ENERGY variety was retained as the most potent commercial variety that would eventually be used as a nutraceutical. The E063 variety was further considered as an alkaloid-rich control given the highest inhibitory effect of its alkaloidic fraction.

### Lupin alkaloids are more potent than non-alkaloid compounds across isolates and nematode species

To further characterize lupin seed inhibitory effect on parasitic nematodes, total seed extracts were fractionated into alkaloidic and non-alkaloidic fractions for both E063 and ENERGY varieties. Alkaloids accounted for 3.30% of the E063 seed mass whereas the alkaloid-poor ENERGY seed alkaloid content amounted 0.043% of the seed mass.

The larval migration assay performed with each extract revealed that alkaloids of both varieties significantly inhibited the larval migration in comparison to negative control across drug-resistance status or nematode species (Fig. 2, supplementary Table 4). Non-alkaloid compounds from ENERGY had no effect on larval migration (12.5% ± 0.06% inhibition difference relative to control, *t*_*39*_ = −2.15, *P* = 0.38), whereas the E063 non-alkaloidic fraction was as potent as the alkaloid fraction against susceptible *H. contortus* larvae (Fig. 2a, supplementary Table 4). E063 total seed extract could inhibit the susceptible *H. contortus* L3 with the same magnitude as the 10µM levamisole solution (99.3% ± 0.06% and 82.8% ± 0.06% mean inhibition for levamisole and E063 total extract respectively after accounting for migration plate effect, *z-score* = −2.84, *P* = 0.09).

**Figure 2.**
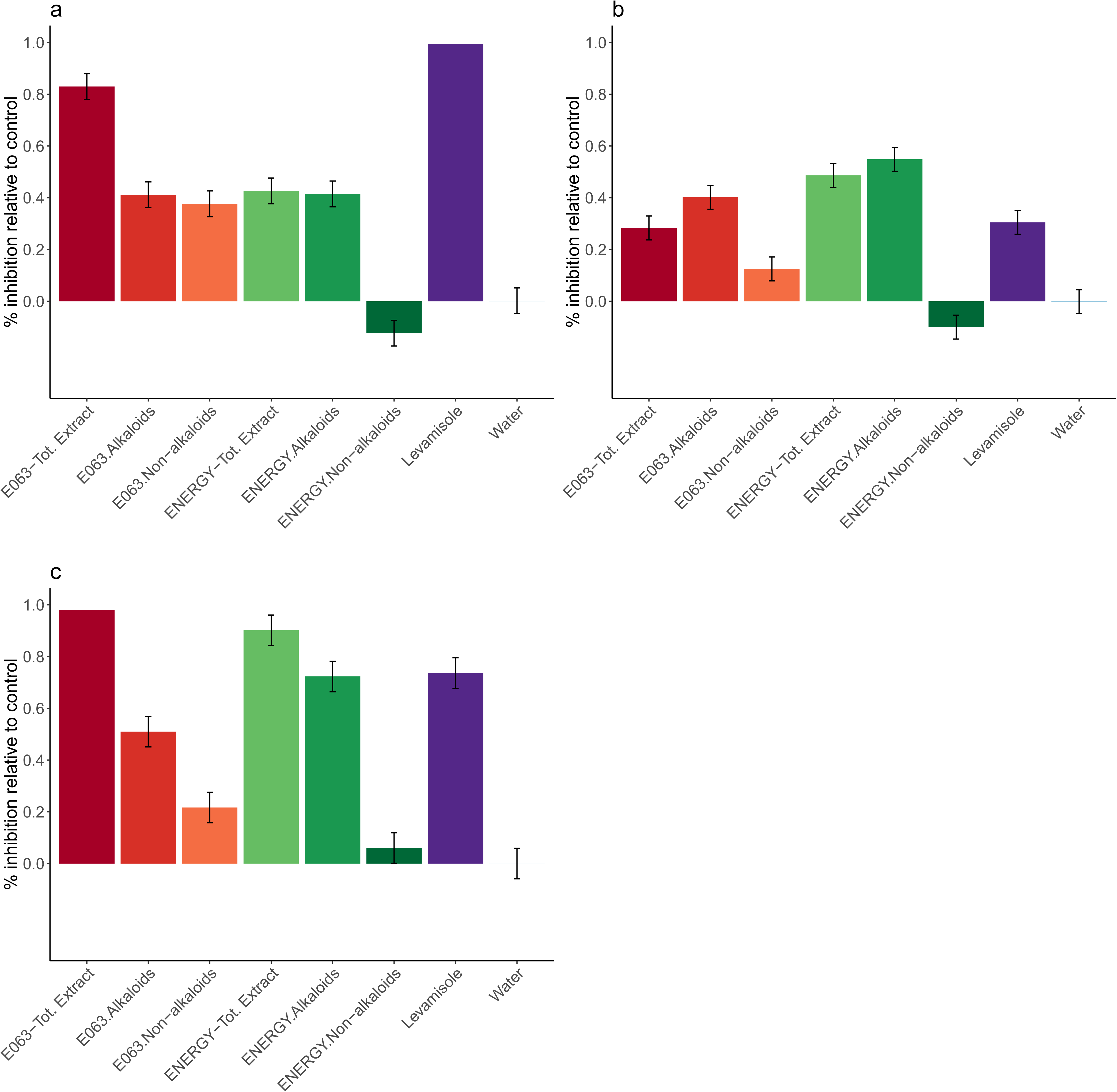
Comparative inhibitory potential of ENERGY and E063 lupin seed total extract, alkaloidic and non-alkaloidic fractions. Picture depicts observed inhibitory effects of lupine seed extracts (expressed as the percentage of observed migration relative to negative control) on drug susceptible (a), multidrug resistant *H. contortus* (b) and susceptible *T. circumcincta* (c) infective larvae migration after exposure to different solutions. “Tot. Extract” stands for total seed extract. Levamisole (10 µM) and water were used as positive and negative controls respectively, and lyophilized extracts were used at a concentration of 5mg/mL. Each condition was run across six replicates.

Alkaloids recovered from ENERGY were significantly more potent than levamisole at inhibiting the migration of the resistant isolate (24.3% ± 0.07% inhibition difference, *z-score* = 3.72, *P* = 5.1 x 10^−3^, Fig. 2b, supplementary Table 4).

Lupin seed total extracts and alkaloid fractions also significantly inhibited *Teladorsagia circumcincta* infective larvae (Fig. 2c, supplementary Table 4).

Altogether these results confirmed the potential of ENERGY and E063 alkaloids as potent anthelmintics against major trichostrongylid of small ruminants and their ability to control multidrug-resistant *H. contortus* isolate.

### Alkaloids of ENERGY and E063 lupin varieties are different and do not exert the same anthelmintic activity

To investigate the difference in the anthelmintic effects exerted by E063 and ENERGY alkaloids, chromatographic analyses were performed on their respective alkaloid fractions (Fig. 3a). Lupanine was the most abundant alkaloid in both cases (21.5% and 49.5% for ENERGY and E063 respectively, supplementary Table 5). ENERGY alkaloids exhibited a more complex alkaloid profile (five major alkaloids identified) than E063 alkaloids (Fig. 3a).

**Figure 3.**
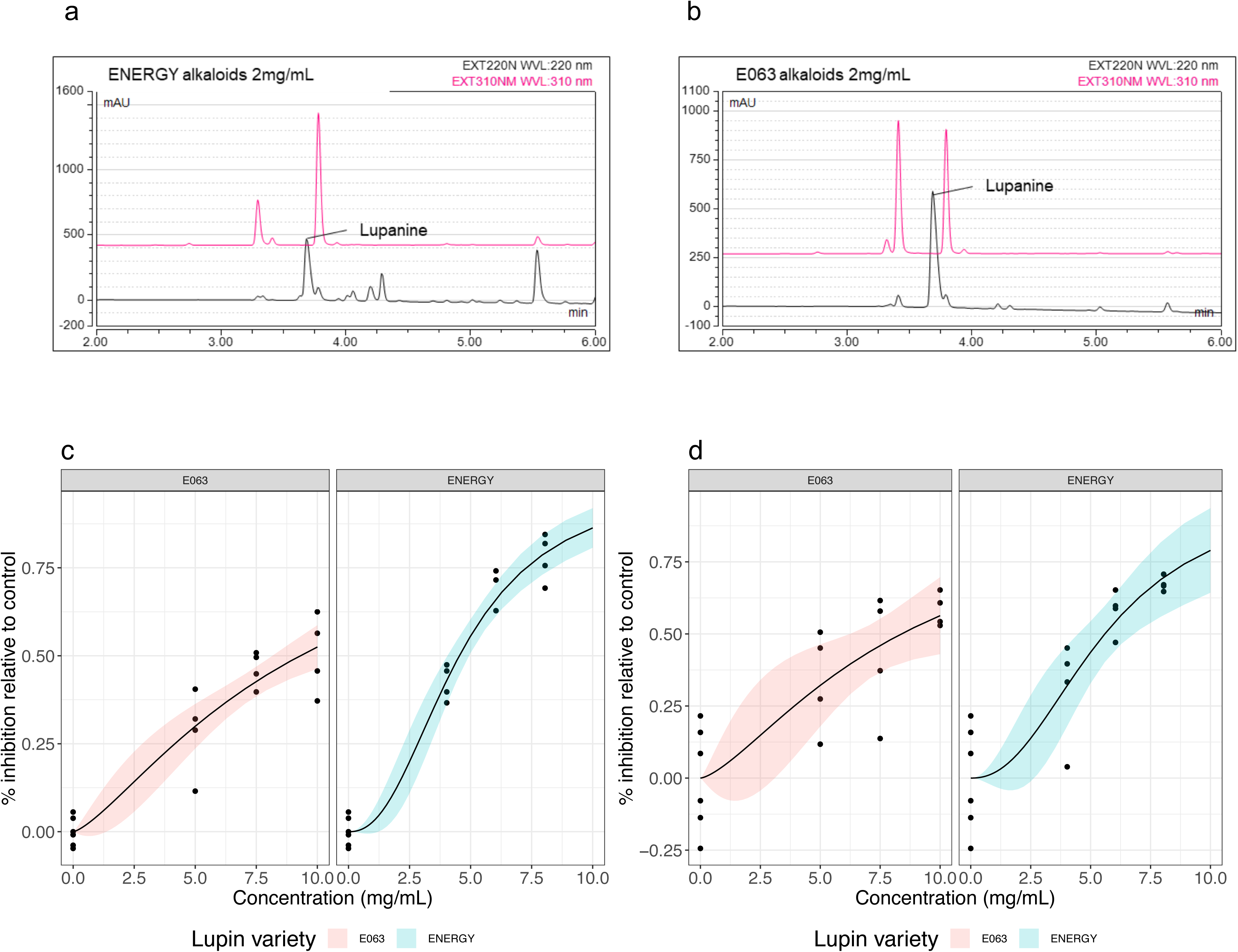
ENERGY and E063 alkaloids differ in their content and anthelmintic effect. HPLC profiles measured at 220 nm (black) and 310 nm (pink) reveal a more diverse alkaloidic content (spikes in profile) in ENERGY (a) than in E063 (b). Bottom panels depict the inhibited fraction of larvae relative to control for alkaloid concentration ranging between 0 and 10 mg/mL, for a drug susceptible (c) and a multidrug resistant (d) isolates. Solid line stands for the fitted log-logistic regression curve and shaded area indicates 95% confidence interval (red for E063 and blue for ENERGY).

Importantly, the qualitative differences in alkaloid profiles between varieties were also associated with contrasted inhibitory effects: estimated ENERGY alkaloids IC_50_ were at least 2 mg/mL lower than for E063 alkaloids across *H. contortus* isolates (differences in IC_50_ between varieties of 4.76 mg/mL, *P*<10^−4^ and 2.79 mg/mL, *P* = 0.03 for the drug-susceptible and resistant isolate respectively; Fig. 3b, supplementary Table 6).

To further delineate the inhibitory effect of alkaloids, a concentration response assay of lupanine, *i.e.* the major alkaloid, was implemented against the susceptible *H. contortus* isolate (Supplementary Fig. 1). While infective larvae responded in a dose-response fashion after levamisole exposure (IC_50_ = 1.61 µM), the lupanine-induced inhibition increased in a linear fashion with lupanine concentration, was highly variable and did inhibit less than 25% of the control migration (Supplementary Fig. 1).

These observations would favour a more potent alkaloid mixture in the ENERGY seeds than in the E063 variety, but lupanine alone could not explain the observed anthelmintic activity.

### Identification of alkaloidic compounds and evaluation of their anthelmintic activity

As lupanine, the most abundant lupin alkaloid, was not underpinning the observed anthelmintic effects of ENERGY alkaloids extract, a Centrifugal Partition Chromatography (CPC) pH zone refining process was used to identify active compounds (supplementary Fig. 2, 3). Ten simplified fractions were obtained (supplementary Fig. 2, 3). Initial screening of their potencies against *H. contortus* by a larval migration inhibition assay revealed that four fractions (7 to 10) were highly active (supplementary Table 7). These alkaloid-only fractions could inhibit between 61.2 ± 5.2% and 93.6% ± 3.7% of larval migration of both drug-susceptible and multidrug-resistant isolates (supplementary Table 7). HPLC analyses indicated that lupanine was not found in any of these fractions but in fraction 5, the potency of which was more limited, *i.e.* 20% and 42.3% inhibition at 2.5 mg/mL for the susceptible and multidrug-resistant isolates respectively, in agreement with observations made for synthetic lupanine (Supplementary Fig. 1).

These four most active fractions (7 to 10) were then purified by semi-preparative HPLC (supplementary table 5) which yielded six pure alkaloids (compounds **1** to **6**; Fig. 4). Because of the limited amount of compound available, their anthelmintic activity was tested using an Automated Larval Migration Assay (ALMA) at concentrations ranging between 250 and 150 µg/mL (Fig. 4). Among the six tested compounds, compounds **5** and **6** did not significantly inhibit drug-susceptible *H. contortus* larval migration (*P* = 0.2 and 0.5, *W* = 2 and 4 respectively; Fig. 4). Larval migration after exposure to other compounds was significantly inhibited falling to 53.6% ± 3.42% s.d. (compounds **1**) to 76% ± 9.68% s.d. (compounds **4**) of control larval migration (*P* = 0.05, *W* = 0, Fig. 4). Compound **1** was also a significant inhibitor of multidrug-resistant larvae which reduced their migration to 62.4% ± 2.88% s.d. (Fig. 4).

**Figure 4.**
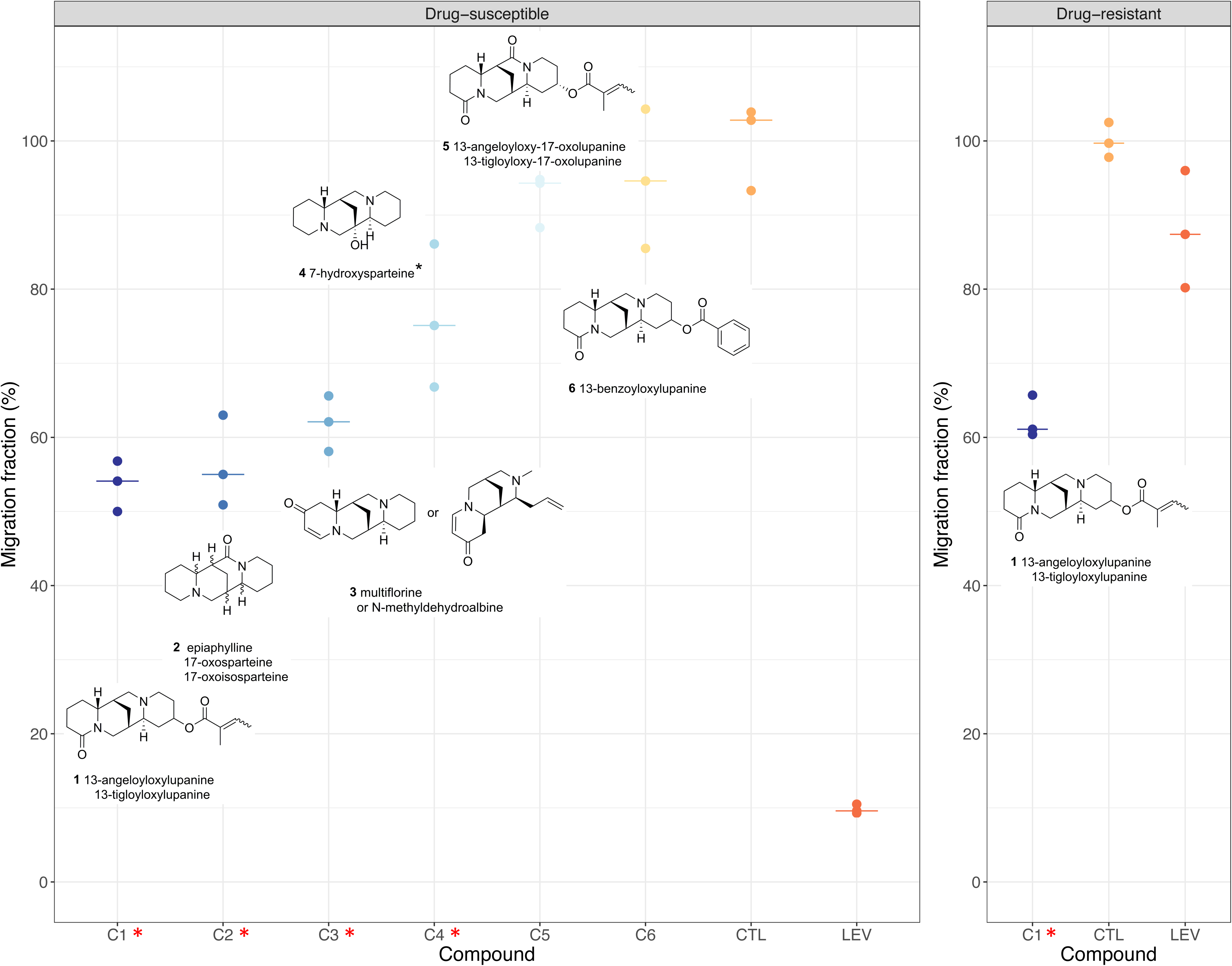
Identified alkaloids and their anthelmintic activity against *H. contortus* infective larvae. Plotted are the results of the Automated Larval Migration Assay expressed as a percentage of drug-susceptible or drug-resistant *H. contortus* larvae migrating (relative to negative control; CTL; water) after exposure to six alkaloidic compounds (**C1** to **C6**; 150 µg/mL for compound **2** and 250 µg/mL else) or appropriate negative (CTL) and positive control (LEV; levamisole 10 µM). Inferred compound structures are provided next to each plot. Bars represent the median migration fraction and red asterisks indicate compound with significant effects.

LC-ESI-MS analysis was performed to identify compound structures by comparison with the litterature. Compound **1** was identified as 13-angeloyloxy or 13-tigloyloxy-lupanine (Fig. 4). Compounds **2** and **4** were identified as sparteine-derivatives, *i.e.* an oxidated analogue of sparteine (epiaphylline, 17-oxosparteine or 17-oxoisosparteine) and 7-hydroxysparteine respectively (Fig. 4). Compound **3** structure was compatible with multiflorane or N-Methyldehydroalbine (Fig. 4). Analytical chemistry combined with the recently developed ALMA unravelled lupanine- and sparteine-derivatives as promising novel chemotherapeutics underpinning the anthelmintic potential of ENERGY seed extracts.

### Electrophysiology reveals an antagonistic mode of action of lupin alkaloids on nematode nicotinic acetylcholine receptors

To elucidate the mechanisms underpinning lupin alkaloids anthelmintic activity, we investigated their interactions with AChRs. We expressed recombinant AChRs, *i.e H. contortus* and *Caenorhabditis elegans* levamisole receptors (Hco-L-AChR-1, Cel-L-AChR respectively) and the nicotine-sensitive *C. elegans* receptor (Cel-N-AChR) in *Xenopus laevis* oocytes and perfused them with ENERGY and E063 extracts as well as lupanine, the most abundant alkaloid lupin extracts (Fig. 5, supplementary Table 8).

**Figure 5.**
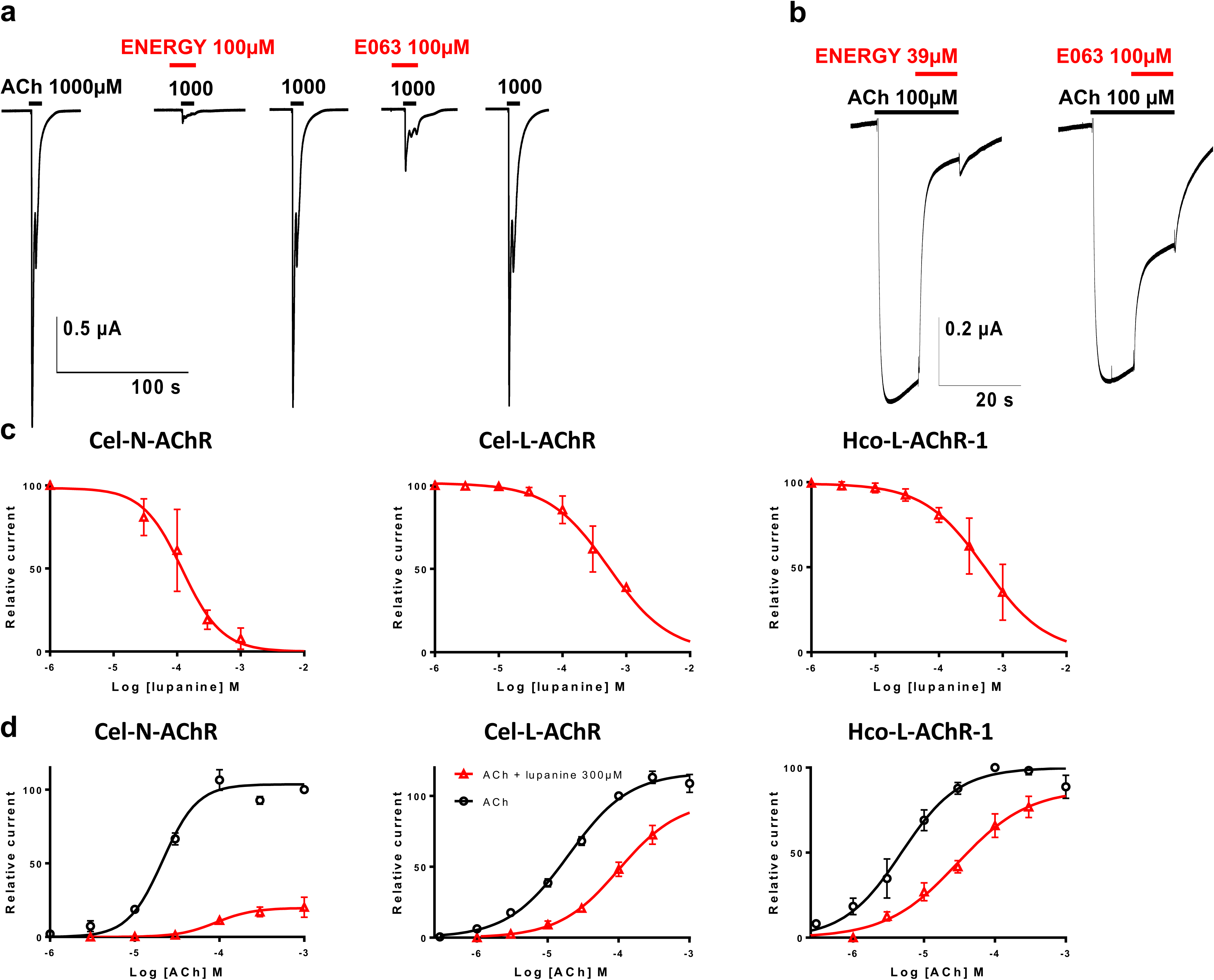
Inhibitory effect of lupin seed extracts from ENERGY and E063 varieties and lupanine on nematode acetylcholine receptor (AChR) subtypes expressed in *Xenopus laevis* oocytes. Figure shows representative recording traces from the *Caenorhabditis elegans* nicotine-sensitive AChR (a) and the *H. contortus* levamisole-sensitive AChR (b) after exposure to acetylcholine (ACh) alone or in the presence of ENERGY and E063 extracts. Concentrations (μM) are indicated above each trace. Bars indicate the time period of the drug application (ACh in black and lupin alkaloids in red). Panel c depicts lupanine concentration-response inhibition curves obtained on Cel-N-AChR (left), Cel-L-AChR (middle) and Hco-L-AChR-1 (right). Panel d displays acetylcholine concentration-response curves either alone (in black), or with 300 µM lupanine (in red) on Cel-N-AChR (left), Cel-L-AChR (middle) and Hco-L-AChR-1 (right).

The application of lupin seed extracts to the oocytes expressing either Hco-L-AChR-1, Cel-L-AChR or Cel-N-AChR never induced any currents thus ruling out a direct agonist effect. However, pre-incubation of *X. laevis* oocytes with ENERGY and E063 extracts significantly blocked the subsequent acetylcholine-elicited currents for every receptor subtype (Fig. 5a, b). The inhibition of ACh-induced current was significantly higher with ENERGY alkaloids than with E063 alkaloids for Hco-L-AChR-1 (91 ± 4% s.d. versus 65 ± 9.6% s.d., *P* = 0.03, paired Wilcoxon’s test = 0; n = 6; Fig. 5b). The magnitude of this effect was more reduced for Cel-N-AChR (87.5 ± 11.2% s.d. versus 64.3 ± 23.7% s.d., *P* = 0.06 paired Wilcoxon’s test; n = 4; Fig. 5a). These antagonistic effects were completely reversible after wash (Fig. 5a).

Because of their limited respective amounts, it was not possible to undertake both ALMA and electrophysiology measurements with alkaloid compounds **1** to **4**. Lupanine is the main alkaloid in both investigated lupin varieties. Despite its lack of direct anthelmintic effect, it was considered further for electrophysiology study to gather insights on the mode of action of its derivatives. Concentration-response assay demonstrated that Cel-N-AChR was more sensitive to the lupanine inhibitory effect on ACh-elicited currents (IC_50_ = 116.5 ± 9.7 µM; Fig. 5c, Supplementary Fig. 3 and supplementary Table 8) whereas a similar inhibitory potential of this alkaloid was observed on levamisole- and nicotine-sensitive receptors (IC_50_ = 539.9 ± 90.2 µM and 548.8 ± 64.1 µM for Hco-L-AChR-1 and Cel-L-AChR respectively; Fig. 5c, Supplementary Fig. 3 and supplementary Table 8). Lupanine-mediated antagonism was further investigated by comparing ACh concentration-response in the absence and in the presence of 300 µM lupanine (Fig. 5d, supplementary Table 8). This assay yielded 3.71-to 5.95-fold increase in ACh EC_50_ values for Hco-L-AChR-1 (EC_50[ACh]_ = 4.8 ± 0.5 µM versus EC_50[ACh+ 300µMLup]_ = 28.6 ± 8.4 µM, *P* = 0.005, *t* = −2.83) and for Cel-N-AChR (EC_50[ACh]_ = 21.7 ± 0.9 µM versus EC_50[ACh+300µM Lup]_ = 80.6 ± 27.5 µM; *P* = 0.034, *t* = −2.14). The same tendency was observed for Cel-N-AChR but was not significant (EC_50[ACh]_ = 19.6 ± 1.7 µM versus EC_50[ACh+300µMLup]_ = 99.1 ± 47.8 µM; *P* = 0.1, *t* = −1.66; Fig. 5d). This variation in acetylcholine affinity across all three receptors is a typical feature of competitive antagonism. Finally, we observed an 81% drop-off in the maximum amplitude of ACh-elicited currents for the Cel-N-AChR whereas this maximal current was less affected (38% diminution) for levamisole-sensitive receptors (Fig. 5d). This would be in favour of a non-competitive antagonism.

Altogether these results suggested that ENERGY potency is mediated by an antagonistic mode of action of its alkaloid content. Electrophysiology also revealed striking differences between the sensitivity of nicotine-sensitive AChR subtypes to lupanine that may underpin the inhibitory potential of lupin extracts against the multidrug-resistant isolate.

### *In vivo* test on growing sheep and dairy goats

The last step of our study was to establish whether ENERGY crude seeds could be used as a nutraceutical. Homogeneous groups of growing sheep and dairy goats (supplementary Table 9) fed with lupin were subjected to experimental infection and their performances compared to respective controls fed with their classical diet (supplementary Table 10). Quantity of lupin in diets (250g/day and 450g/day for ewes and goats respectively, supplementary table 10) was maximized but constrained to feed animals with diets balanced for protein and energy requirements. As a result of the infection by *H. contortus*, Faecal Egg Count (FEC, Fig. 6a, b) increased throughout the experiment and reached maximal values at 30 days post-infection (dpi) whereas parasitic-mediated blood losses inflected haematocrit values (1.83% ± 0.45 and 1.5% ± 0.44 differences relatively to 18 dpi for ewes and for goats across conditions, Fig. 6c, d).

**Figure 6.**
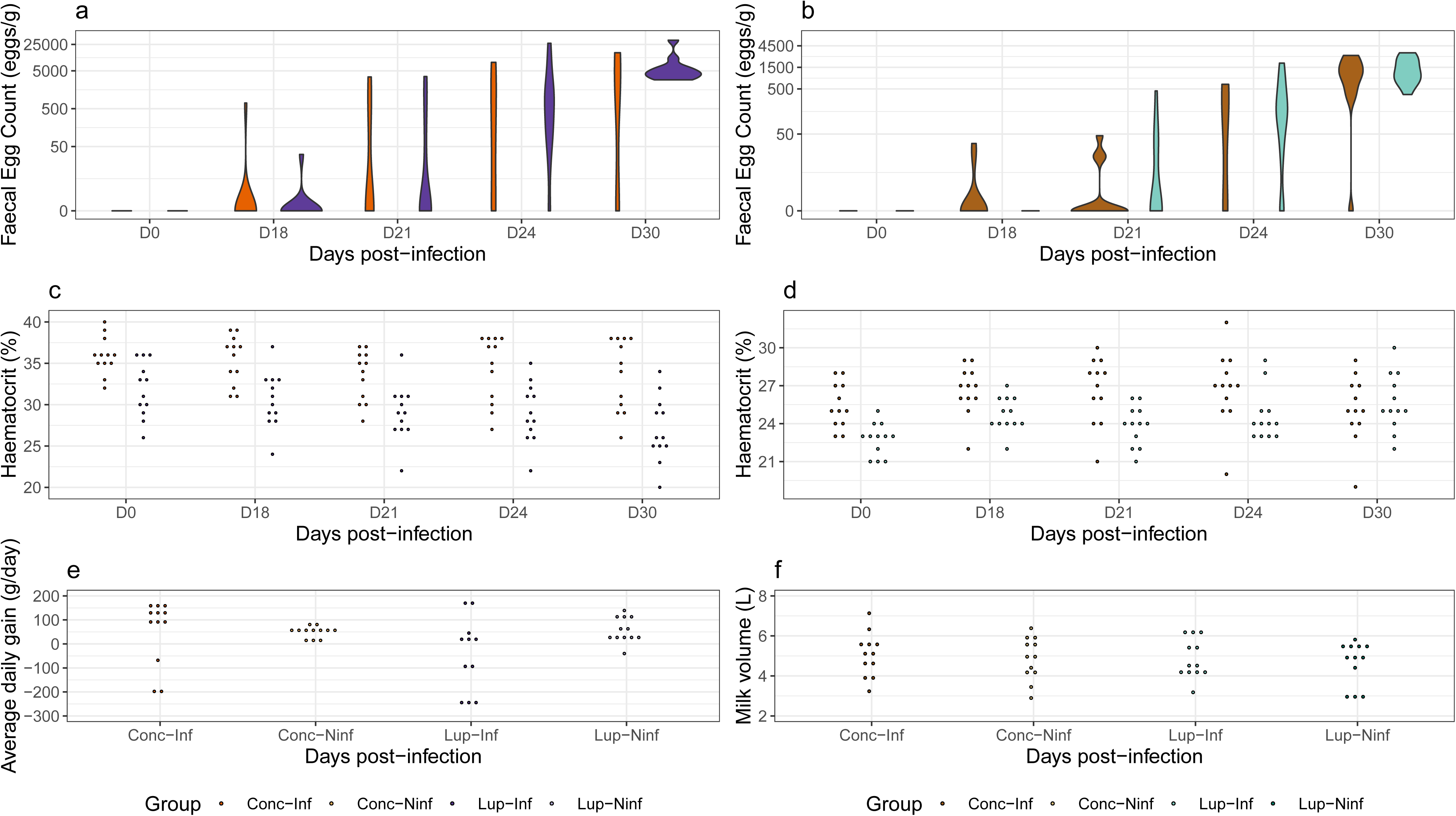
*In vivo* test of lupin energy seed on sheep and goats challenged with *H. contortus*. Figure shows measured Faecal Egg Counts (a, b), haematocrit (c, d) and production traits (e, f) for growing ewes (a, c, e) and dairy goats (b, d, f). Average daily gain is the growth difference between 30 and 0 dpi scaled by the number of days. Milk volume is the sum of three recorded milk yields at 0, 21 and 30 dpi. Dots are coloured by experimental groups (Lup-Inf: lupin-fed and infected; Lup-Ninf: lupin-fed and not infected; Conc-Inf: concentrate-fed and infected; Conc-Ninf: concentrate-fed and not infected)

FEC did not show significant differences between lupin-fed and concentrate-fed animals suggesting lupin seed did not affect *H. contortus* within its host (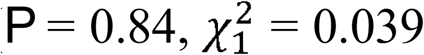 for ewes; 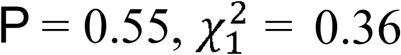 for goats; Fig. 5a, b). On the contrary, ewes fed with lupin had higher FEC values at 30 dpi than their concentrate-fed counterparts (116 eggs/g difference between groups, *P* = 2 x 10^−3^, *t*_*66*_ = 3.2; Fig. 6a). Also, lupin seeds did not delay the onset of egg excretion in sheep or goats and did not alter the larval development rate measured that both remained similar between lupin-fed and others in ewes and goats (supplementary Table 11).

Ewes resilience was not different between both groups as similar haematocrits were found throughout the challenge (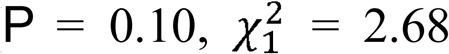 Fig. 6b). However, goats exhibited a slightly different pattern: the overall trend in blood losses was not significantly different between diets (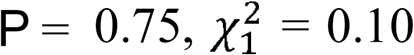 Fig. 6c) but blood losses were significantly reduced in goats fed with lupin at 30 dpi (2.41% ± 0.63 difference in haematocrit between both groups, *P* = 3 x 10^−4^, *t*_*66*_ = 3.86; Fig. 6c). This would suggest an increased resilience in dairy goats fed with lupin seeds. But this increased resilience was not observed for production traits. Growth rate deteriorated in infected ewes fed with lupin seeds (−122 g/day 46.1, P = 0.01, *t*_*3*_ = −2.644; Fig. 6e) and goat milk yield remained unchanged between groups throughout the experimental challenge (*P* = 0.83, *F*_*3,44*_ = 0.3; Fig. 6e).

Despite the promising compounds harboured in ENERGY lupin seeds, these observations suggest that commercial lupin seed cannot be used to control major *H. contortus* infection in growing ewes at the dose tested in this setting. Still, it may contribute to dampen blood losses suffered by infected goats.

## Discussion

We provide first line of evidence to support the anthelmintic properties of lupin seed extracts against parasitic nematodes. The seed alkaloid content was mediating an anthelmintic effect which was conserved across drug resistance status and trichostrongylid species. Analytical chemistry revealed four lupanine- and sparteine-derivatives as the most promising novel anthelmintic leads. This anthelmintic effect could result from both competitive and non-competitive inhibition of parasitic nematode acetylcholine receptors as suggested by electrophysiology data. Furthermore, commercial lupin variety conferred increased resilience to *H. contortus* infected dairy goats but had no direct effect on faecal egg excretion or larval development rate.

Lupin seed alkaloids could hence contribute to the development of novel compounds able to break resistance as observed inhibitory effects were shared across trichostrongylid species. This finding is in line with previous works in helminths of veterinary or medical importance^31-34^ or plant-parasitic nematodes^35,36^ that also evidenced the anthelmintic potential of plant extracted alkaloids. Our results also suggested that lupin extracts would control multidrug-resistant field isolate, which is an important consideration given the threat to sustainable control impeded by such isolates worldwide, including Europe^37^.

Electrophysiology of nematode recombinant receptors in *Xenopus* oocytes contributed to resolve some of the molecular bases underpinning the lupin seed extract inhibitory effects. While lupanine exerted similar antagonistic effect against cholinergic receptors, concentration-response curves suggested contrasted pharmacological interactions of lupanine with levamisole- and nicotine-sensitive receptor subtypes. Of note, EC_50_ shifts observed for levamisole-sensitive receptors would suggest a reversible competitive antagonist mode of action, whereas currents measured on Cel-N-AChR suggests that lupanine acts as a non-competitive antagonist. As these cholinergic receptors control muscle cell contractions at neuromuscular junctions, the resistance breaking effect of lupine seed extracts may be mediated by the simultaneous action upon several receptor subtypes. This hypothesis underscores the importance of nAChRs for the development of novel anthelmintic compounds^38^ and highlights the need to investigate the putative modulatory effect of lupin alkaloids on other nematode ligand-gated ion channel subtypes.

Resolving the contribution of AChRs antagonism to the observed anthelmintic effects would require additional experiments that are beyond the scope of this study. For such experiment, the knock-down of acetylcholine receptor expression by RNA interference^39^ in parasitic larvae before further exposure to lupin alkaloids could be a method of choice. However, our results indicated that the E063 enriched alkaloidic fraction was not necessarily exerting as much inhibition as the total seed extract and that significant larval migration inhibition was associated with the non-alkaloid seed extract in that case. This suggests that lupin seed extracts may contain chemicals which would have additional effects on parasitic larvae. It may be speculated that some non-alkaloid compounds, like flavonoids, could prevent alkaloid metabolism by the parasite as this is the case with ivermectin^40^. This would explain the relatively low effect observed for lupanine on larval migration that contrasts with its clear antagonistic effect on cholinergic receptors.

Results from the *in vivo* experiment were also promising regarding the nutraceutical potential of lupin seeds. Unlike alkaloid-rich lupin varieties, commercial lupin seeds contain only traces of alkaloids^41^ that are responsible for seed bitterness ^23^, hence making lupin seeds palatable enough for livestock species^18,19^ as observed in our trial on ewes and goats. Despite the lack of beneficial effects in ewes and the similar egg excretion levels monitored between diets, blood losses were lessened in goats at the end of their challenge. Lupin seeds could hence contribute to better sustain *H. contortus*-infected goat welfare; its omega-6 fatty acids content might underpin a beneficial pro-clotting effect as seen in hens^42^. However, the residual alkaloid content harboured in commercial lupin seeds does not seem to suffice to exert any direct anthelmintic effect, especially in ewes which received less seeds in this trial beause of their reduced ingestion capacity. Additional developments are hence required to deliver lupin-based solution to breeders. Selection of lupin varieties with higher alkaloid contents added to livestock diets could be an option as well as the combination of lupin seeds with tannin-rich forages.

In conclusion, we provided first evidence of lupin seed anthelmintic properties against major trichostrongylid of ruminants and identified lupanine- and sparteine-derivatives as the likely active compounds. Our results also suggested that alkaloid compounds act as antagonist of parasitic cholinergic receptors. While lupin seed did not contribute to reduce parasite egg excretion in experimentally challenged ruminants, it increased goat resilience. Lupin seeds may hence serve as a nutraceutical to increase the sustainability of ruminant farming by both contributing to limit anthelmintic usage and supplying protein requirements. More importantly, alkaloid compounds harboured in commercial lupin seed provide a working basis for the development of novel anthelmintic compounds able to break drug resistance in the field.

## Materials and methods

### Ethics statement

Animal experiment was approved by the Centre Val de Loire Ethics committee (Licence number APAFIS#2224-2015100917282331 v2) and carried out according to EU regulation on animal experiment (2010/63/UE).

### Lupin species and varieties

An initial screening for anthelmintic activity was performed on 11 populations of lupin seeds, encompassing four different species, *i.e. L. albus, L. angustifolius, L. luteus* and *L. mutabilis* that exhibited either high or low alkaloid content (Supplementary Table 1). Alkaloid-rich seeds are not commercially available and were provided by the national biobank for proteaginous plants (CRB Protéagineux, INRA, Dijon, France) as well as the LANG172 and LL049 varieties. Other commercial varieties with low alkaloid content, namely CLOVIS, ENERGY, and ORUS were provided by Jouffray-Drillaud (Lusignan, France).

### Lupin seed extract preparation

To prepare lupin seed aqueous extracts used in the initial screening, seeds were crushed into powder and incubated within water (1/5 w/w) during 48h at 20 ± 2 °C. Aqueous extract was then filtered on cotton mesh before being frozen and lyophilised. For parasitological tests, lyophilised extracts were diluted in water at 5 mg/mL and filtered through 0.2 µm mesh to increase solution transparency and prevent the obstruction of the migration mesh, thereby preventing any interaction with the larval migration assay.

### Alkaloid extraction, fractionation and identification

Lupin alkaloids extraction protocol was adapted from previous work^43^. Lupin seeds were crushed and extracted five times with HCl 0.5 M (1/6, w/w), before pH adjustment to 12 with 40% NaOH and three subsequent extraction with CH_2_Cl_2_ (6:1, v/v). Resulting organic phases were pooled, dried with MgSO_4_ and concentrated under vacuum before three additional extractions with HCl 0.05M (3:1, v/v). Residual traces of CH_2_Cl_2_ were eliminated by evaporation under vacuum, and the resulting aqueous alkaloidic extract was freeze dried.

Compositional insights on the alkaloidic fractions was subsequently inferred by ultra-high-performance thin layer chromatography (UHPLC). For UHPLC, mobile phases were solvent A 0.1% TFA in water and solvent B acetonitrile. Gradient was set by raising acetonitrile content from 0% to 100% in 15 min and maintained for 2 min, with a flow rate of 0.8 mL.min-1, and oven temperature set at 40°C. Chromatogram was monitored at 205, 220 and 305 nm. Calibration curve was derived using 5 µL of 200, 400, 600, 800 and 1,000 µg/mL lupanine perchlorate (Innosil, Poznán, Poland) solutions. Lupin aqueous maceration or alkaloids extract were solubilized in water to obtain a final concentration between 1 and 20 mg/mL as requested.

To identify the alkaloidic compounds harboured by Energy seeds, crude alkaloids extract (1.9 g) obtained from 3.2 kg of seeds were fractionated using Centrifugal Partition Chromatography (CPC) in pHzone refining mode with a biphasic solvent system in descending mode (Methyl terbutyl ether/n-Butanol/water, 23:32:45, v/v/v). Triethylamine (29.4 mM) was used as a retainer in organic stationary phase, and HCl (3.3 mM) was used as a displacer in aqueous mobile phase. Chromatographic parameters were set as follows: flow rate 6 mL.min-1; rotation speed 1400 rpm; back pressure 40 bars; stationary phase retention 70%. Resulting fractions were purified using semi-preparative HPLC with the same UHPLC chain (See supplementary material). Isolated alkaloids were then analysed using mass spectrometry (LC-ESI-MS) as described elsewhere^44^. Compounds structure were proposed according to retention time combined with m/z and fragmentation profile by comparison with literature^45^.

### Larval migration inhibition assay test (LMIA)

Parasite material was maintained by the INRA animal parasite nematode collection^46^. Two *H. contortus* isolates with contrasted resistance status, *i.e.* susceptible or resistant to every licensed anthelmintic compound in France (benzimidazoles, levamisole, macrocyclic lactones) were used. To establish whether lupin effect was conserved across GIN species, a fully susceptible French *Teladorsagia circumcincta* isolate was also used for *in vitro* testing. For every test, infective larvae were exsheathed with 0.5% hypochlorite solution and rinsed thrice with water before subsequent incubation in 5 mg/mL solutions of either crude maceration, alkaloidic fraction or non-alkaloidic fraction, levamisole (10 µM) or water, for 18h at 20°C. At the end of the incubation period, 100 L3 were deposited on migration plates with 20 µm mesh at 38°C for 2h as described elsewhere^47^. Typically, for each variety of the initial screening on 11 varieties, three replicates were made and normalized by six control observations. Dose-response assays were run with six control replicates and four replicates by concentration tested (0, 5, 7.5 and 10 mg/mL for E063 alkaloids and 0, 2, 4, 6 and 8 mg/mL for ENERGY alkaloids).

### Automated Larval Migration Assay (ALMA) for the isolated alkaloid from energy seeds

Because of the scarcity of fractionated alkaloidic compounds (2.7 to 12.8 mg), their anthelmintic effect on infective *H. contortus* larvae was measured using ALMA. This technic relies on larval auto-fluorescence generated through ultraviolet excitation^38,39^. Every alkaloidic compound was incubated for 4 hours at 20°C with 7,500 *H. contortus* infective larvae each. Larvae were then passed through a 20⁏μm sieve and left for a 60 s stabilization time before migration into the spectrofluorimeter cuvette. Cumulative fluorescence resulting from larval migration into the cuvette and correlating the total number of larvae^39^ was recorded during 5⁏min. Each set of experiment was performed in triplicate at a concentration of 250 µg/mL but for compound **2** (150 µg/mL). Levamisole 10µM was used as a positive control.

### *Xenopus laevis* oocyte electrophysiology

The lupanine and alkaloid extracts from lupin seeds were perfused on defolliculated *X. laevis* oocytes (Ecocyte Bioscience, Dortmund, Germany) expressing either the nicotine-sensitive nAChR subtype from *Caenorhabditis elegans* (Cel-N-nAChR) encoded by *acr-16*^48^ or the levamisole-sensitive nAChR subtype 1 (Hco-L-nAChR1) from *H. contortus* encoded by *unc-29.1, unc-38, unc-63* and *acr-8*^26^ respectively. Expression of recombinant nAChRs was achieved by microinjection of cRNAs into the oocyte cytoplasm and two-electrode voltage-clamp recordings were carried out as previously described^26^.

In any case, acetylcholine was applied first to check for the presence of functional nAChRs and for current normalization in presence of lupanine or alkaloid extracts. Antagonist modulation of acetylcholine-elicited current was determined by pre-application of lupanine or alkaloid extracts for 10 seconds, followed by their co-application in the presence of acetylcholine.

Data were collected and analysed using pCLAMP 10.4 package (Molecular Devices, UK).

### *In vivo* testing

Any nutraceutical should show in vitro anthelmintic activity, be palatable and exert its effect in the infected hosts^15^. To test the last two properties, 48 female Romane ewe and 48 Alpine dairy goats were allocated to four experimental groups corresponding to possible meal composition and infection status (**Lup-Inf**: lupin-fed and infected; **Lup-Ninf**: lupin-fed and not infected; **Conc-Inf**: concentrate-fed and infected; **Conc-Ninf**: concentrate-fed and not infected). Dietary supplementation with protein can increase host resilience to *H. contortus* infection^49^. Therefore, lupin- and concentrate-diets were designed to be balanced in their total energy and protein content (supplementary Table 10) and to fulfil animal nutritional needs (a growth rate of 150 g/day for ewes and a 3.1 kg/day milk production for goats)^50^. Lupin- and concentrate-diets were similar in energy (around 1,580 and 3,600 kcal NEL, for ewes and goats respectively), proteins (around 13% and 14% CP, for ewes and goats respectively) and fibers (around 20% and 22% CF, for ewes and goats respectively).

The experimental infection was run indoor and was set up to mimic field chronic infection. It consisted in a first infection with 5,000 *H. contortus* larvae that lasted 30 days before animals were treated (ivermectin for ewes, Oramec^®^, 2.5 mL/10 kg bodyweight, Merial, France; fenbendazole for goats, Panacur 2.5% NDV, 10 mg/kg, MSD, France) and left indoors for 15 days. A second challenge then took place with 5,000 *H. contortus* and blood and faecal samples were taken just before infection, at 18-, 21-, 24- and 30-days post-infection (dpi) onwards to measure haematocrit and Faecal Egg Count (FEC). Ewes were weighted at every time point and goat milk production was recorded (morning milking) at 0, 21 and 30 dpi. To evaluate any putative effect of lupin on larval development, faecal samples (10g) were collected at 30 dpi from the two most infected individuals, mixed together and allocated to six culture batches incubated for 12 days at 25°C and 60% humidity). Infective third stage larvae were then enumerated and the ratio between observed and expected (derived from the total number of eggs used for incubation) number of larvae, *i.e.* larval development rate was derived.

### Statistical tests

The inhibitory potential of each variety was computed as:

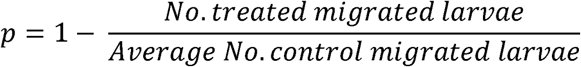

This proportion was subsequently regressed upon variety and parasitic isolate, simultaneously accounting for the migration plate effect fitted as a random effect with the lme function of the R software v3.5^51^. Model diagnostics was run to check for residual normality and detect putative outliers that were discarded accordingly. Post-hoc comparisons accounting for multiple comparisons between tested effect levels were performed using a Tukey’s test implemented with the *multcomp* package v1.4-8^52^.

Dose-response data were modelled as log-logistic regression with two, *i.e.* the curve slope and the associated median excitatory (EC50) or inhibitory (IC50) concentration using the *drc* package v3.0-1^53^. A third parameter was added for electrophysiology data modelling to account for differences in maximal response values.

Linear regression of FEC and haematocrit upon experimental group, day post infection and their interaction considered as fixed effects and individual as a random effect was implemented with the *lme* function of the R v3.5 software^50^. To account for group heterogeneity in haematocrit at the beginning of the 2^nd^ infection (FEC were 0 for every individual at 0 dpi), basal haematocrit level measured at 0 dpi was used as covariates. Average daily gain was computed for every ewe as the difference between weights at 30 and 0 dpi divided by the number of days, and recorded milk production volumes were summed for each goat as a proxy for their total volume. Both traits were regressed upon experimental group accounting for an individual random effect. Every measured trait but FEC was normally distributed (assessed by Shapiro-Wilk’s test values of 0.85 and 0.81 for average daily gain and transformed FEC, and above 0.96 for other traits). FEC were thus applied a fourth-root transformation to correct for normality departure.

Average prepatent period length and larval development rate were compared between diets within species using a Wilcoxon’s test.

Unless stated otherwise, average estimates are reported with their standard error (s.e.m.).

## Data availability

Data have been provided as supplementary information. Raw data are available upon requests to the corresponding author.

## Acknowledgements

This project has received the grateful support of the Centre-Val de Loire region (Lupinema project, grant number 2014-00091669). Alkaloid-rich seeds were provided by the INRA CRB Protéagineux and nematodes were maintained by the BRC4Env Animal Parasitic Nematodes collection, the environmental resources pillar of the AgroBRC-RARe research infrastructure. Authors are grateful to N. Harzic for providing commercial lupin seeds.

## Authors’ contribution

GS, LBD, CLC, CEG, and CN designed the project. OD, LBD, ITK performed analytical chemistry. OD, FG, HF and GS implemented *in vitro* tests. CLC, OD, EC and NP performed and analyzed electrophysiology measurements. CK and JC harvested parasite material and performed larval development assay. GS, CH, AM and FB designed and supervised the *in vivo* experiment. CA and GS took care of sampling sheep and goats and analysed collected data. JBMR harvested lupin seeds. GS, LBD and CLC drafted the manuscript.

## References

1. Crellen, T. et al. Reduced Efficacy of Praziquantel Against Schistosoma mansoni Is Associated With Multiple Rounds of Mass Drug Administration. Clin Infect Dis 63, 1151–1159, doi:10.1093/cid/ciw506 (2016).

2. Geurden, T. et al. Anthelmintic resistance to ivermectin and moxidectin in gastrointestinal nematodes of cattle in Europe. Int J Parasitol Drugs Drug Resist 5, 163–171, doi:10.1016/j.ijpddr.2015.08.001 (2015).

3. Matthews, J. B. Anthelmintic resistance in equine nematodes. Int J Parasitol Drugs Drug Resist 4, 310–315, doi:10.1016/j.ijpddr.2014.10.003 (2014).

4. Paraud, C. et al. Cross-resistance to moxidectin and ivermectin on a meat sheep farm in France. Vet Parasitol 226, 88–92, doi:10.1016/j.vetpar.2016.06.033 (2016).

5. Dalys, G. B. D. & Collaborators, H. Global, regional, and national disability-adjusted life-years (DALYs) for 333 diseases and injuries and healthy life expectancy (HALE) for 195 countries and territories, 1990-2016: a systematic analysis for the Global Burden of Disease Study 2016. Lancet 390, 1260–1344, doi:10.1016/S0140- 6736(17)32130-X (2017).

6. Kaplan, R. M. & Vidyashankar, A. N. An inconvenient truth: Global worming and anthelmintic resistance. Vet Parasitol 186, 70–78, doi:10.1016/j.vetpar.2011.11.048 (2012).

7. Sackett, D. H. P.; Abott, K.; Jephcott S.; Barber M. Assessing the economic cost of endemic disease on the profitability of Australian beef cattle and sheep producers. Animal health and welfare report to Meat and Livestock Australia. (Meat and Livestock Australia Ltd, Sidney, 2006).

8. Kaminsky, R. et al. A new class of anthelmintics effective against drug-resistant nematodes. Nature 452, 176– 180, doi:10.1038/nature06722 (2008).

9. Scott, I. et al. Lack of efficacy of monepantel against Teladorsagia circumcincta and Trichostrongylus colubriformis. Vet Parasitol 198, 166–171, doi:10.1016/j.vetpar.2013.07.037 (2013).

10. Hewitson, J. P. & Maizels, R. M. Vaccination against helminth parasite infections. Expert Rev Vaccines 13, 473–487, doi:10.1586/14760584.2014.893195 (2014).

11. Sallé, G. et al. Transcriptomic profiling of nematode parasites surviving vaccine exposure. Int J Parasitol, doi:10.1016/j.ijpara.2018.01.004 (2018).

12. Riggio, V. et al. A joint analysis to identify loci underlying variation in nematode resistance in three European sheep populations. J Anim Breed Genet 131, 426–436, doi:10.1111/jbg.12071 (2014).

13. Assenza, F. et al. Genetic parameters for growth and faecal worm egg count following Haemonchus contortus experimental infestations using pedigree and molecular information. Genet Sel Evol 46, 13, doi:10.1186/1297- 9686-46-13 (2014).

14. Kenyon, F. et al. The role of targeted selective treatments in the development of refugia-based approaches to the control of gastrointestinal nematodes of small ruminants. Vet Parasitol 164, 3–11, doi:10.1016/j.vetpar.2009.04.015 (2009).

15. Hoste, H. et al. Tannin containing legumes as a model for nutraceuticals against digestive parasites in livestock. Vet Parasitol 212, 5–17, doi:10.1016/j.vetpar.2015.06.026 (2015).

16. Williams, A. R. et al. Anthelmintic activity of trans-cinnamaldehyde and A- and B-type proanthocyanidins derived from cinnamon (Cinnamomum verum). Sci Rep 5, 14791, doi:10.1038/srep14791 (2015).

17. Cronk, Q., Ojeda, I. & Pennington, R. T. Legume comparative genomics: progress in phylogenetics and phylogenomics. Curr Opin Plant Biol 9, 99–103, doi:10.1016/j.pbi.2006.01.011 (2006).

18. May, M. G. et al. Lupins (Lupinus albus) as a protein supplement for lactating Holstein dairy cows. J Dairy Sci 76, 2682–2691, doi:10.3168/jds.S0022-0302(93)77604-3 (1993).

19. Watkins, B. A. & Mirosh, L. W. White lupin as a protein source for layers. Poult Sci 66, 1798–1806 (1987).

20. Cabello-Hurtado, F. et al. Proteomics for exploiting diversity of lupin seed storage proteins and their use as nutraceuticals for health and welfare. J Proteomics 143, 57–68, doi:10.1016/j.jprot.2016.03.026 (2016).

21. Jimenez-Lopez, J. C. et al. Narrow-Leafed Lupin (Lupinus angustifolius) beta1-and beta6-Conglutin Proteins Exhibit Antifungal Activity, Protecting Plants against Necrotrophic Pathogen Induced Damage from Sclerotinia sclerotiorum and Phytophthora nicotianae. Front Plant Sci 7, 1856, doi:10.3389/fpls.2016.01856 (2016).

22. Villate, L., Morin, E., Demangeat, G., Van Helden, M. & Esmenjaud, D. Control of Xiphinema index populations by fallow plants under greenhouse and field conditions. Phytopathology 102, 627–634, doi:10.1094/PHYTO-01-12-0007 (2012).

23. Green, B. T., Welch, K. D., Panter, K. E. & Lee, S. T. Plant toxins that affect nicotinic acetylcholine receptors: a review. Chem Res Toxicol 26, 1129–1138, doi:10.1021/tx400166f (2013).

24. Yovo, K. et al. Comparative pharmacological study of sparteine and its ketonic derivative lupanine from seeds of Lupinus albus. Planta Med 50, 420–424, doi:10.1055/s-2007-969753 (1984).

25. Beech, R. N. & Neveu, C. The evolution of pentameric ligand-gated ion-channels and the changing family of anthelmintic drug targets. Parasitology 142, 303–317, doi:10.1017/S003118201400170X (2015).

26. Boulin, T. et al. Functional reconstitution of Haemonchus contortus acetylcholine receptors in Xenopus oocytes provides mechanistic insights into levamisole resistance. Br J Pharmacol 164, 1421–1432, doi:10.1111/j.1476-5381.2011.01420.x (2011).

27. Robertson, S. J., Pennington, A. J., Evans, A. M. & Martin, R. J. The action of pyrantel as an agonist and an open channel blocker at acetylcholine receptors in isolated Ascaris suum muscle vesicles. Eur J Pharmacol 271, 273–282 (1994).

28. Baur, R., Beech, R., Sigel, E. & Rufener, L. Monepantel irreversibly binds to and opens Haemonchus contortus MPTL-1 and Caenorhabditis elegans ACR-20 receptors. Mol Pharmacol 87, 96–102, doi:10.1124/mol.114.095653 (2015).

29. Blanchflower, S. E., Banks, R. M., Everett, J. R., Manger, B. R. & Reading, C. New paraherquamide antibiotics with anthelmintic activity. J Antibiot (Tokyo) 44, 492–497 (1991).

30. Zinser, E. W. et al. Anthelmintic paraherquamides are cholinergic antagonists in gastrointestinal nematodes and mammals. J. vet. Pharmacol. Therap. 25, 241–250, doi:10.1046/j.1365-2885.2002.00423.x (2002).

31. Wangchuk, P., Giacomin, P. R., Pearson, M. S., Smout, M. J. & Loukas, A. Identification of lead chemotherapeutic agents from medicinal plants against blood flukes and whipworms. Sci Rep 6, 32101, doi:10.1038/srep32101 (2016).

32. Satou, T. et al. Assay of nematocidal activity of isoquinoline alkaloids using third-stage larvae of Strongyloides ratti and S. venezuelensis. Vet Parasitol 104, 131–138 (2002).

33. Perrett, S. & Whitfield, P. J. Atanine (3-dimethylallyl-4-methoxy-2-quinolone), an alkaloid with anthelmintic activity from the Chinese medicinal plant, Evodia rutaecarpa. Planta Med 61, 276–278, doi:10.1055/s-2006- 958073 (1995).

34. Ayers, S. et al. Anthelmintic activity of aporphine alkaloids from Cissampelos capensis. Planta Med 73, 296– 297, doi:10.1055/s-2007-967124 (2007).

35. Wen, Y. et al. Nematotoxicity of drupacine and a Cephalotaxus alkaloid preparation against the plant-parasitic nematodes Meloidogyne incognita and Bursaphelenchus xylophilus. Pest Manag Sci 69, 1026–1033, doi:10.1002/ps.3548 (2013).

36. Thoden, T. C., Boppre, M. & Hallmann, J. Effects of pyrrolizidine alkaloids on the performance of plant-parasitic and free-living nematodes. Pest Manag Sci 65, 823–830, doi:10.1002/ps.1764 (2009).

37. Blake, N. & Coles, G. Flock cull due to anthelmintic-resistant nematodes. Vet Rec 161, 36 (2007).

38. Charvet, C. L., Guégnard, F., Courtot, E., Cortet J. & Neveu, C. Nicotine-sensitive acetylcholine receptors are relevant pharmacological targets for the control of multidrug resistant parasitic nematodes. International Journal for Parasitology: Drug and Drug Resistance 8, 540–549, doi:10.1016/j.ijpddr.2018.11.003 (2018).

39. Blanchard, A. et al. Deciphering the molecular determinants of cholinergic anthelmintic sensitivity in nematodes: When novel functional validation approaches highlight major differences between the model Caenorhabditis elegans and parasitic species. PLoS Pathog 14, e1006996, doi:10.1371/journal.ppat.1006996 (2018).

40. Bartley, D. J. et al. P-glycoprotein interfering agents potentiate ivermectin susceptibility in ivermectin sensitive and resistant isolates of Teladorsagia circumcincta and Haemonchus contortus. Parasitology 136, 1081–1088, doi:10.1017/S0031182009990345 (2009).

41. Boschin, G., Annicchiarico, P., Resta, D., D’Agostina, A. & Arnoldi, A. Quinolizidine alkaloids in seeds of lupin genotypes of different origins. J Agric Food Chem 56, 3657–3663, doi:10.1021/jf7037218 (2008).

42. Yeh, E., Wood, R. D., Leeson, S. & Squires, E. J. Effect of dietary omega-3 and omega-6 fatty acids on clotting activities of Factor V, VII and X in fatty liver haemorrhagic syndrome-susceptible laying hens. Br Poult Sci 50, 382–392, doi:10.1080/00071660902942767 (2009).

43. Ganzera, M., Kruger, A. & Wink, M. Determination of quinolizidine alkaloids in different Lupinus species by NACE using UV and MS detection. J Pharm Biomed Anal 53, 1231–1235, doi:10.1016/j.jpba.2010.05.030 (2010).

44. Mulder, P. P. et al. Transfer of pyrrolizidine alkaloids from various herbs to eggs and meat in laying hens. Food Addit Contam Part A Chem Anal Control Expo Risk Assess 33, 1826–1839, doi:10.1080/19440049.2016.1241430 (2016).

45. Wink, M., Meisser, C. & Witte, L. Patterns of quinolizidine alkaloids in 56 species of the genus Lupinus. Phytochem. 38, 139–153 (1995).

46. Mougin, C. et al. BRC4Env, a network of Biological Resource Centres for research in environmental and agricultural sciences. Environ Sci Pollut Res Int, doi:10.1007/s11356-018-1973-7 (2018).

47. Kotze, A. C., Le Jambre, L. F. & O’Grady, J. A modified larval migration assay for detection of resistance to macrocyclic lactones in Haemonchus contortus, and drug screening with Trichostrongylidae parasites. Vet Parasitol 137, 294–305, doi:10.1016/j.vetpar.2006.01.017 (2006).

48. Boulin, T. et al. Eight genes are required for functional reconstitution of the Caenorhabditis elegans levamisole-sensitive acetylcholine receptor. Proc Natl Acad Sci U S A 105, 18590–18595, doi:10.1073/pnas.0806933105 (2008).

49. Coop, R. L. & Kyriazakis, I. Influence of host nutrition on the development and consequences of nematode parasitism in ruminants. Trends Parasitol 17, 325–330 (2001).

50. Abad, P. et al. Genome sequence of the metazoan plant-parasitic nematode Meloidogyne incognita. Nat Biotechnol 26, 909–915, doi:10.1038/nbt.1482 (2008).

51. R: A Language and Environment for Statistical Computing (R Foundation for Statistical Computing, Vienna, 2016).

52. Hothorn, T., Bretz, F. & Westfall, P. Simultaneous inference in general parametric models. Biom J 50, 346– 363, doi:10.1002/bimj.200810425 (2008).

53. Ritz, C., Baty, F., Streibig, J. C. & Gerhard, D. Dose-Response Analysis Using R. PLoS One 10, e0146021, doi:10.1371/journal.pone.0146021 (2015).

