## Supplementary material for "Lupin (*Lupinus sp.*) seeds exert anthelmintic activity associated with their alkaloid content"

### Supplementary information

**Supplementary Table 1. Description of the 11 lupin varieties considered in the initial screening for nematicide effect**

| Species | Variety | Origin | Alkaloid content |
|---|---|---|---|
| *L. albus* | E063 | Spain | High |
| *L. albus* | EGY014 | Egypt | High |
| *L. albus* | EGY100 | Egypt | High |
| *L. angustifolius* | LANG061 | France | High |
| *L. luteus* | LL151 | Portugal | High |
| *L. mutabilis* | LM261 | Peru | High |
| *L. albus* | CLOVIS* | France | Low |
| *L. albus* | ENERGY* | France | Low |
| *L. albus* | ORUS* | France | Low |
| *L. angustifolius* | LANG172 | Australia | Low |
| *L. luteus* | LL049 | Germany | Low |

*, commercially available varieties in Europe

**Supplementary Table 2. Measured inhibitory effect of 11 lupin variety extracts on *Haemonchus contortus* infective larvae**

| Lupin variety | Alkaloid content | *H. contortus* resistance status | Average inhibitory effect | Standard deviation |
|---|---|---|---|---|
| CLOVIS | Low | Multidrug resistant | 61.97% | 5.59% |
| Control | Low | Multidrug resistant | 5.28% | 6.14% |
| E063 | High | Multidrug resistant | 78.10% | 10.03% |
| EGY014 | High | Multidrug resistant | 84.67% | 3.79% |
| EGY100 | High | Multidrug resistant | 81.75% | 3.34% |
| ENERGY | Low | Multidrug resistant | 75.35% | 4.88% |
| LANG061 | High | Multidrug resistant | 70.80% | 12.83% |
| LANG172 | Low | Multidrug resistant | 59.86% | 11.76% |
| LL049 | Low | Multidrug resistant | 73.24% | 5.32% |
| LL151 | High | Multidrug resistant | 63.50% | 22.37% |
| LM261 | High | Multidrug resistant | 83.94% | 9.12% |
| ORUS | Low | Multidrug resistant | 58.45% | 5.32% |
| CLOVIS | Low | Fully susceptible | 33.94% | 12.00% |
| Control | Low | Fully susceptible | 2.52% | 4.18% |
| E063 | High | Fully susceptible | 53.15% | 9.96% |
| EGY014 | High | Fully susceptible | 95.95% | 2.70% |
| EGY100 | High | Fully susceptible | 66.22% | 4.68% |
| ENERGY | Low | Fully susceptible | 40.37% | 5.73% |
| LANG061 | High | Fully susceptible | 69.37% | 6.80% |
| LANG172 | Low | Fully susceptible | 8.41% | 9.56% |
| LL049 | Low | Fully susceptible | 45.41% | 3.18% |
| LL151 | High | Fully susceptible | 50.90% | 12.26% |
| LM261 | High | Fully susceptible | 92.79% | 0.78% |
| ORUS | Low | Fully susceptible | 27.98% | 7.83% |

**Supplementary Table 3. Measured inhibitory effect of the alkaloidic fractions recovered from alkaloidic-rich varieties on *Haemonchus contortus* infective larvae**

| Lupin variety | *H. contortus* resistance status | Average inhibitory effect | Standard deviation |
|---|---|---|---|
| Water control | Multidrug resistant | 0.00% | 12.90% |
| E063 | Multidrug resistant | 81.99% | 3.58% |
| EGY014 | Multidrug resistant | 72.99% | 8.65% |
| EGY100 | Multidrug resistant | 73.46% | 8.33% |
| LANG061 | Multidrug resistant | 69.67% | 5.75% |
| LL151 | Multidrug resistant | 59.24% | 10.67% |
| LM261 | Multidrug resistant | 78.20% | 6.41% |
| Water control | Fully susceptible | 0.00% | 10.77% |
| E063 | Fully susceptible | 77.59% | 3.95% |
| EGY014 | Fully susceptible | 61.49% | 7.53% |
| EGY100 | Fully susceptible | 73.28% | 3.95% |
| LANG061 | Fully susceptible | 54.31% | 9.00% |
| LL151 | Fully susceptible | 52.01% | 2.77% |
| LM261 | Fully susceptible | 56.03% | 2.28% |

**Supplementary Table 4. Compared inhibitory potentials of alkaloidic and non-alkaloidic fractions extracted from the ENERGY and E063 varieties against drug susceptible and -resistant *H.contortus* and drug-susceptible *T. circumcincta***

| | ***H. contortus* - Drug susceptible** | | ***H. contortus* - Multidrug resistant** | | ***T. teladorsagia* - Susceptible** | |
|---|---|---|---|---|---|---|
| | Average | Std. | Average | Std. | mean | Std. |
| Water | 0.17% | 5.42% | -0.17% | 8.42% | 0.00% | 13.49% |
| Levamisole | 99.50% | 1.22% | 30.50% | 6.38% | 73.70% | 6.68% |
| E063-Total Extract | 83.00% | 4.90% | 28.33% | 7.28% | 98.00% | 2.68% |
| E063.Alkaloids | 41.17% | 13.53% | 40.17% | 9.06% | 51.00% | 9.14% |
| E063.Non-alkaloids | 37.67% | 7.79% | 12.50% | 6.41% | 21.70% | 8.19% |
| ENERGY-Total Extract | 42.67% | 8.24% | 48.67% | 15.55% | 90.20% | 8.95% |
| ENERGY.Alkaloids | 41.50% | 16.56% | 54.83% | 9.06% | 72.30% | 9.79% |
| ENERGY.Non-alkaloids | -12.33% | 15.63% | -10.00% | 20.34% | 6.00% | 18.81% |

Average inhibitory effect and corresponding standard deviations are presented for each every tested condition across *H.contortus* isolates and a susceptible *T. circumcincta* isolate. Each condition was run in triplicate. Levamisole was used at 10 µM and lyophilized extracts were used at a concentration of 5mg/mL.

**Supplementary table 5. Lupanine base content of ENERGY and E063 alkaloid fractions quantified by UHPLC**

| **Lupin variety** | **Lupanine base (µg)** | **Extract injected (µg)** | **Percentage of Lupanine base (%)** |
|---|---|---|---|
| ENERGY | 2.15 ± 0.02 | 10 | 21.5 ± 0.2 |
| E063 | 2.97 ± 0.01 | 6 | 49.5 ± 0.2 |

**Supplementary Table 6. Median inhibitory concentrations ($IC_{50}$) estimated for ENERGY and E063 alkaloid fractions on *H.contortus* infective larvae**

| | | $IC_{50}$ | Std. | Lower bound | Upper bound |
|---|---|---|---|---|---|
| Multidrug resistant *H. contortus* | E063 | 8.38 | 1.15 | 6.05 | 10.72 |
| | ENERGY | 5.59 | 0.47 | 4.64 | 6.55 |
| Fully susceptible *H. contortus* | E063 | 9.30 | 0.73 | 7.82 | 10.78 |
| | ENERGY | 4.54 | 0.21 | 4.11 | 4.96 |

$IC_{50}$ was estimated from a log-logistic regression. *Std* indicates estimated standard deviation, and lower and upper bound of the 95% confidence interval are provided.

**Supplementary Table 7. Average migration of *H. contortus* larvae after exposure to ENERGY CPC-fractionated alkaloidic extracts**

| Isolate | Fraction | Percentage of migrating larvae | Standard deviation |
|---|---|---|---|
| Multidrug-resistant | Crude alkaloid extract | 43.8 | 11.7 |
| | F1 | 91.7 | 17.4 |
| | F2 | 0.8 | 1.4 |
| | F3 | 2.5 | 2.5 |
| | F4 | 120.7 | 24.5 |
| | F5 | 58.7 | 5.2 |
| | F6 | 58.7 | 20.2 |
| | F7 | 38.8 | 5.2 |
| | F8 | 14.9 | 5.0 |
| | F9 | 22.3 | 5.0 |
| | F10 | 6.6 | 3.8 |
| | Levamisole (10 µM) | 39.7 | 8.9 |
| | Negative control | 100.0 | 19.9 |
| Fully susceptible | Crude alkaloid extract | 49.3 | 2.1 |
| | F1 | 91.4 | 24.7 |
| | F2 | 15.0 | 2.1 |
| | F3 | 5.7 | 4.5 |
| | F4 | 90.0 | 9.3 |
| | F5 | 80.0 | 24.4 |
| | F6 | 74.3 | 5.0 |
| | F7 | 33.6 | 6.2 |
| | F8 | 17.9 | 2.5 |
| | F9 | 15.7 | 1.2 |
| | F10 | 6.4 | 3.7 |
| | Levamisole (10 µM) | 12.1 | 5.4 |
| | Negative control | 100.0 | 12.6 |

Results of a Larval Migration Inhibition Assay are provided for ten fractions obtained after CPC-fractionation of ENERGY alkaloidic extracts and corresponding crude extract. Percentages and standard deviations were estimated from three replicates. Water was used a negative control. Fractions were tested at a concentration of 2.5 mg/mL, but fractions 1 and 8 (0.625 mg/mL).

**Supplementary Table 8. Estimated median excitatory concentrations ($EC_{50}$) for acetylcholine in absence or presence of lupanine and inhibitory concentrations ($IC_{50}$) of lupanine on nematode cholinergic receptors**

| | Cel-N-AChR | Cel-L-AChR | Hco-L-AChR-1 |
|---|---|---|---|
| ***Acetylcholine alone*** | | | |
| $EC_{50}$ (μM) | 21.7 ±0.9 | 19.6 ±1.7 | 4.8 ±0.5 |
| n | 12 | 11 | 10 |
| ***Acetylcholine with Lupanine (300 μM)*** | | | |
| $EC_{50}$ (μM) | 80.6 ±27.5 | 99.1 ±47.8 | 28.6 ±8.4 |
| n | 5 | 6 | 8 |
| ***Lupanine*** | | | |
| $IC_{50}$ (μM) | 116.5 ±9.7 | 548.8 ±64.1 | 539.9 ±90.2 |
| n | 5 | 5 | 5 |

$EC_{50}$ of acetylcholine and $IC_{50}$ of lupanine and standard deviation are reported for *Caenorhabditis elegans* nicotine- (Cel-N-AChR) and levamisole-sensitive (Cel-L-AChR) receptors, and *H. contortus* levamisole-sensitive AChRs (Hco-L-AChR-1). *n* refers to the number of *X. laevis* oocytes measured in each case.

**Supplementary Table 9. Average production traits and dispersion for considered experimental groups at the start of the experiment**

| Species | Trait | Lup-Inf | Lup-Ninf | Conc-Inf | Conc-Ninf |
|---|---|---|---|---|---|
| Goat | Milking rank | 1.9 ± 1.1 [1;4] | 2.5 ± 1.3 [1;5] | 2.1 ± 1.0 [1;4] | 2.3 ± 1.4 [1;6] |
| | Milking stage (days) | 103 ± 18 [71;129] | 116 ± 16 [84;131] | 102 ± 20 [68;122] | 107 ± 20 [77;128] |
| | Milking volume (L) | 1.8 ± 0.3 [1.4;2.2] | 1.6 ± 0.5 [0.8;2.4] | 2.1 ± 0.4 [1.5;2.6] | 1.9 ± 0.4 [1.4;2.5] |
| Sheep | Age (days) | 124 ± 2.1 [121;129] | 125 ± 2.0 [122;129] | 124 ± 2.3 [121;128] | 123 ± 2.5 [120;129] |
| | Weight (Kg) | 36.1 ± 4.2 [26.8;41.5] | 37.3 ± 3.1 [31.1;42.7] | 36.1 ± 3.3 [32.7;43.6] | 36.4 ± 3.3 [30.6;42.5] |

Summary statistics of ewes and goats production traits are listed for the four considered experimental groups (Lup-Inf: lupin-fed and infected; Lup-Ninf: lupin-fed and not infected; Conc-Inf: concentrate-fed and infected; Conc-Ninf: concentrate-fed and not infected). Statistics are given as "mean ± standard deviation [minimum ; maximum]" and computed from 12 individuals in each case.

**Supplementary Table 10. Diet compositions for considered experimental groups**

| Species | Group | Ingredient | Quantity |
|---|---|---|---|
| Sheep | Conc-Inf/Conc-Ninf | Commercial concentrate | 0.51 |
| | | Hay | 0.66 |
| | Lup-Inf/Lup-Ninf | Lupin | 0.25 |
| | | Barley | 0.17 |
| | | Hay | 0.66 |
| Goat | Conc-Inf/Conc-Ninf | Rapeseed meal | 0.63 |
| | | Commercial concentrate | 0.22 |
| | | Hay | 2 |
| | Lup-Inf/Lup-Ninf | Lupin | 0.45 |
| | | Commercial concentrate | 0.22 |
| | | Hay | 2 |

Ingredients quantity are given in kg of organic matter/individual/day for each dietary type in both species. Lup-Inf: lupin-fed and infected; Lup-Ninf: lupin-fed and not infected; Conc-Inf: concentrate-fed and infected; Conc-Ninf: concentrate-fed and not infected.

**Supplementary Table 11. Prepatent period length and larval development rate measured in infected sheep and goats**

| Species | Infected group | Prepatent period (days) | Larval development rate (%) |
|---|---|---|---|
| Ewe | Lup-Inf | 23.3 ± 2.9 | 53.3 ± 25.5 |
| | Conc-Inf | 24.8 ± 4.3 | 55.2 ± 22.3 |
| Goat | Lup-Inf | 24.2 ± 3.5 | 32.0 ± 15.7 |
| | Conc-Inf | 25.3 ± 4.6 | 34.3 ± 21.9 |

Prepatent period was computed as the average number of days before the onset of egg excretion within group (n = 12). Larval development rate was estimated from 6 replicates. Results are reported as "mean ± standard deviation". Lup-Inf: lupin-fed and infected; Conc-Inf: concentrate-fed and infected.

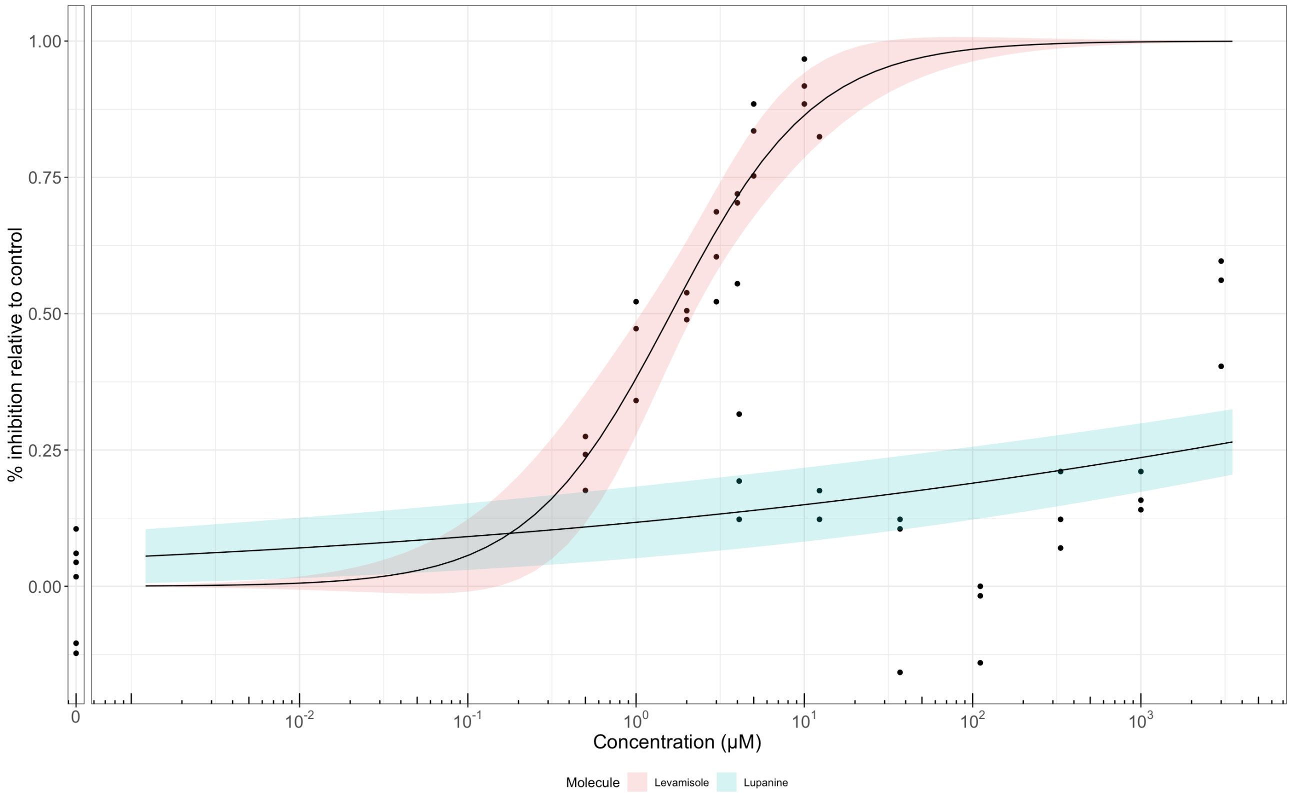

**Figure S1. Concentration-response curves for lupanine and levamisole obtained from larval migration inhibition assay on susceptible *H.contortus* infective larvae**

Plot shows the inhibited fraction of larvae relative to control for Lupanine and levamisole concentration ranging between 0 and 3 mM. Solid line stands for the fitted log-logistic regression curve and shaded area indicates 95% confidence interval.

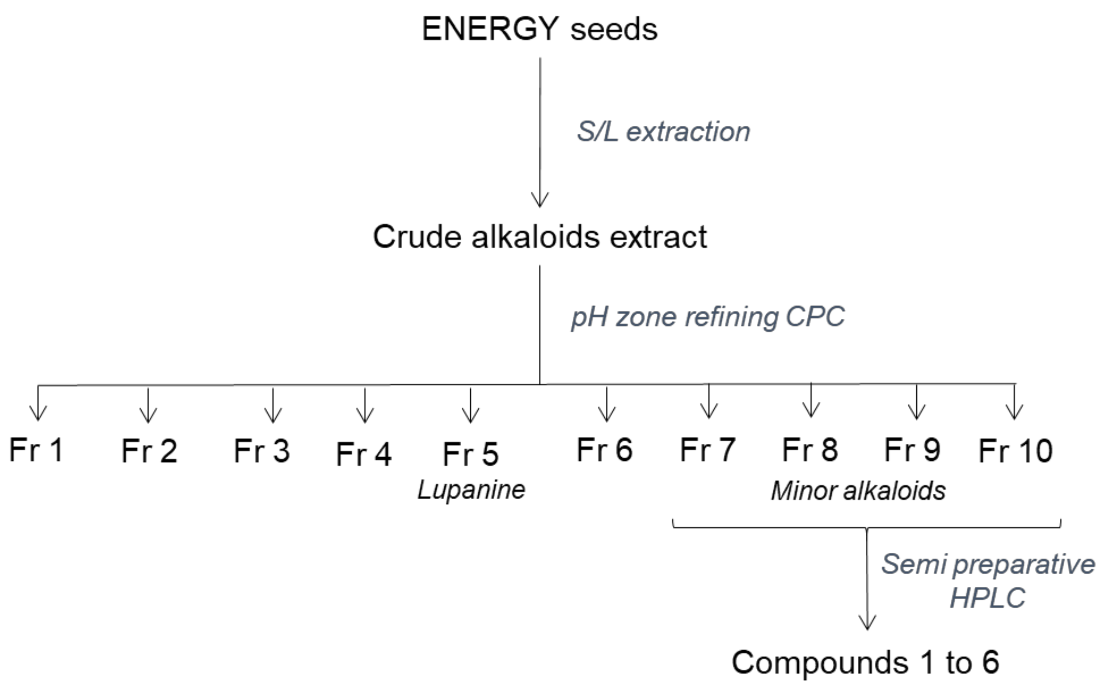

**Figure S2: Alkaloid extraction and identification scheme for the energy lupin variety**

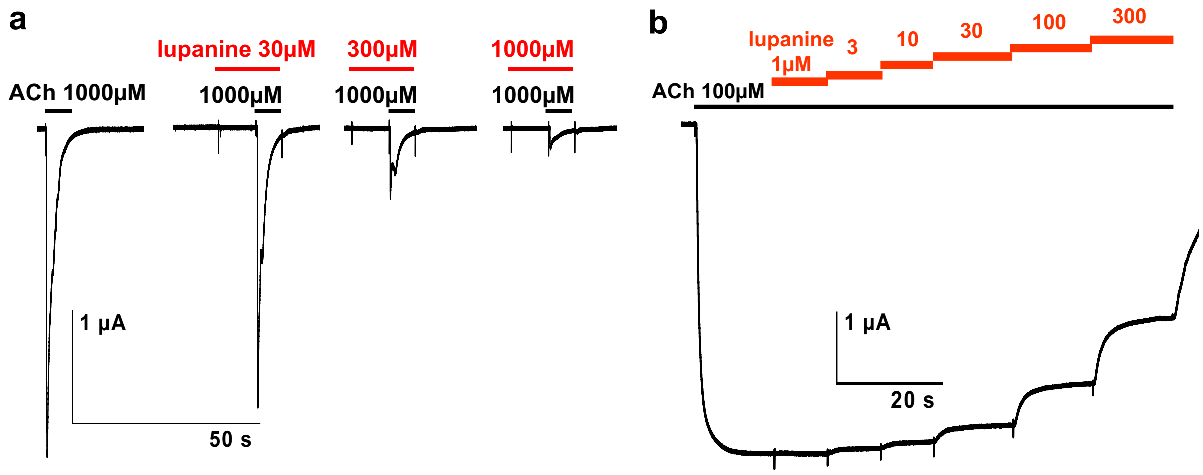

**Figure S3. Inhibitory effect of lupanine on recombinant acetylcholine receptors**

Representative recording traces of ACh response (black bar) in the presence of increasing concentrations of lupanine (red bars) on the *C. elegans* N-AChR (a) and the Hco-L-AChR-1 (b) are plotted.

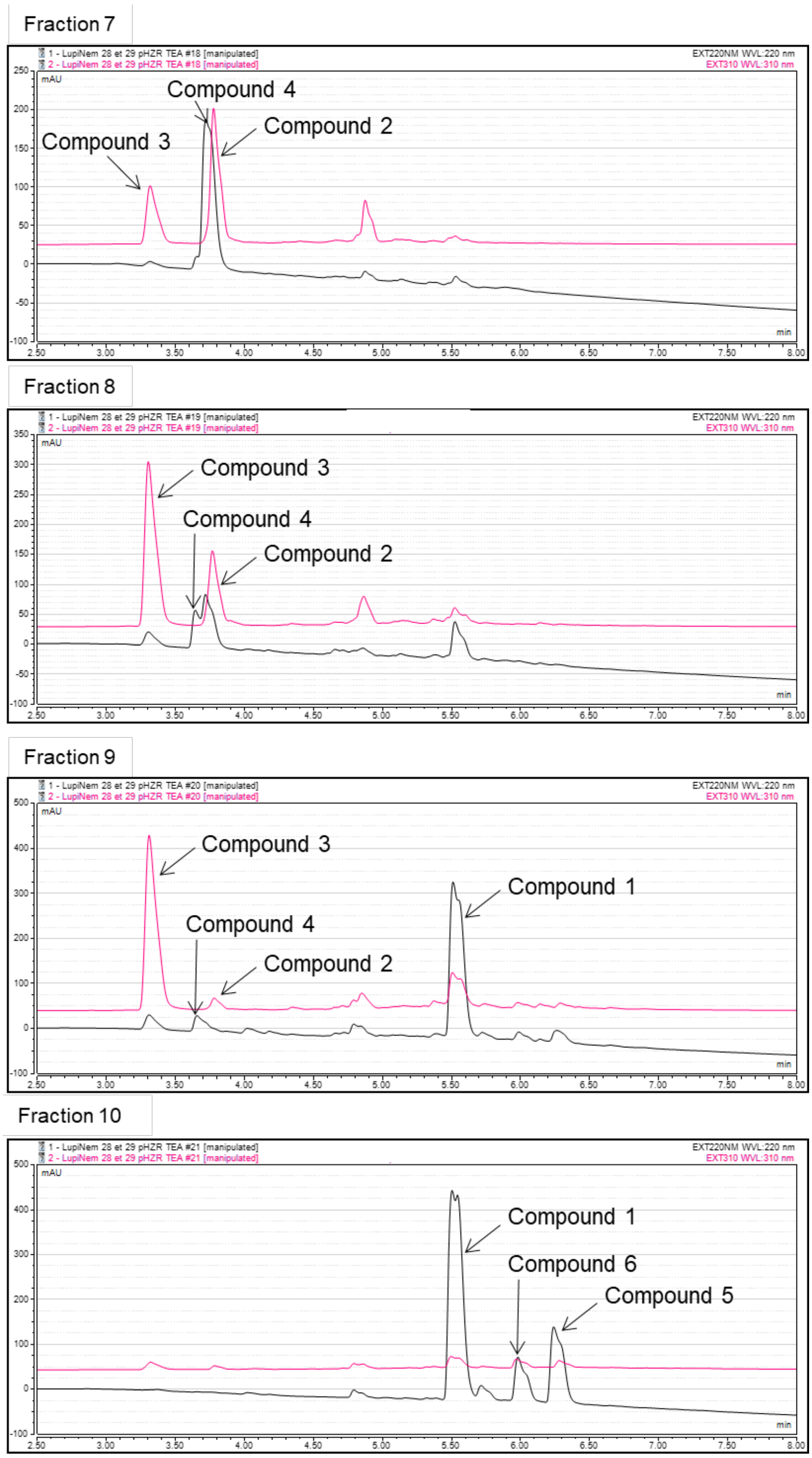

**Figure S5. HPLC Chromatogram of active alkaloids fractions obtained after CPC fractionation**

HPLC profile at 220 nm (black) and 310 nm (pink) of fractions 7 to 10 were represented here.
